# Bacterial suppression of intestinal fungi via activation of human gut γδ T-cells

**DOI:** 10.1101/2025.09.17.676560

**Authors:** Liya Mathew, Sean Carlson, Michael Savage, Claire Pardieu, Megan O’Brien, Ellie Crawley, Paul Harrow, Ann-Kristin Kaune, Edward Devlin, Emily L. Priest, Marilena Crescente, Kimberly Martinod, James Boot, Marco Gasparetto, Klaartje Kok, James O. Lindsay, Jonathan P. Richardson, Julian R. Naglik, Andrew J. Stagg, Matthias Eberl, Siu Kwan Sze, Carol A. Munro, Neil E. McCarthy

## Abstract

Gut symbionts condition mucosal immunity to resist infection by enteropathogens, but the specific microbes and mechanisms involved differ significantly between host species. In higher primates, bacterial metabolite HMB-PP is sensed by a specialized population of Vγ9Vδ2+T-cells, which we now report can potently suppress growth of endogenous fungi in human intestinal organ cultures. In healthy intestine, HMB-PP-stimulated Vδ2+T-cells restricted outgrowth of keystone fungus *Candida albicans* via a mechanism that required IL-22. In contrast, Crohn’s disease (CD) patients with reduced Vδ2+T-cell numbers displayed outgrowth of *C. albicans* strains that readily formed toxin-producing filaments, triggered neutrophil extracellular traps, and induced macrophage IL-1β release *ex vivo*. Genomic and proteomic analysis of the *Candida* isolates suggested increased tissue adhesion of CD-derived strains, which rapidly invaded the gut barrier in an ‘intestine-on-a-chip’ model. These data reveal that bacterial activation of Vδ2+T-cells suppresses fungal pathobionts in human gut via an IL-22-dependent mechanism that is dysregulated in CD.

## Introduction

Commensal microbes condition mucosal immunity to protect against pathogen invasion and limit infection-related tissue damage^1–3^, but the specific species and mechanisms involved differ between host animals and the diverse communities that colonize them^4^. Commensal metabolites have previously been shown to prevent colitis in mice infected with *Helicobacter hepaticus*^5^, and modify the course of *Salmonella typhimurium* infection in rodents^6^ and nematodes^7^, whereas disruption of these mechanisms confers increased risk of mucosal inflammation^8^. However, several major enteropathogens of primates including *Norovirus* and *Shigella* are absent or avirulent in small animal models^4,9,10^, hence the commensal-driven mechanisms that limit these infections in humans are largely unknown.

Pathobiont fungus *Candida albicans* routinely colonizes the human gut^11^, where change in the host environment can stimulate production of tissue-invasive hyphal filaments which secrete candidalysin (CLYS) toxin and damage epithelial barriers. Intestinal *C. albicans* is a potent inducer of both mucosal and systemic Th17 immunity in humans^12^, but does not naturally colonize the mouse gut, which is instead populated by alternative fungal species that stimulate Th2 responses^13^. In immunocompromised patients, *C. albicans* invasion of gut tissue can lead to disseminated infection with mortality >50%, hence the World Health Organization has now identified this species as a critical priority pathogen^14^. Furthermore, recent reports suggest that *C. albicans*-mediated damage to the gut epithelium and resident leukocytes may also play a major role in inflammatory bowel disease (IBD)^15^. Importantly, *C. albicans* pathogenicity can be antagonized by commensal bacteria^16^, which exhibit reduced numbers and species diversity in the inflamed gut^17^, but the host immune mechanisms involved in fungal suppression in human intestine are poorly understood.

At microbe-colonized body sites, commensal signals are rapidly sensed by various types of unconventional lymphocyte^18^, which display marked differences in tissue frequency and cytokine profiles between mice and humans. Mucosal-associated invariant T-cells represent the single largest antigen-specific lineage in the human immune system, but this population is strikingly rare in laboratory mice, which instead feature higher numbers of natural killer T-cells^19,20^. The γδ T-cell compartments of mice and primates are also distinct in key respects, including stark differences in expression of the major antifungal cytokine IL-17^21^, and complete absence of the bacterial metabolite-responsive Vδ2+ population in rodents^22^. In the current report, we observed that Vδ2+T-cell activation by bacterial metabolite (*E*)-4-Hydroxy-3-methyl-but-2-enyl pyrophosphate (HMB-PP) can suppress a range of fungal pathobionts in human gut tissue via an IL-22-dependent mechanism that is dysregulated in Crohn’s disease (CD). Consequently, *C. albicans* strains derived from CD gut biopsy cultures readily formed toxin-producing filaments, triggered release of neutrophil extracellular traps, and induced macrophage IL-1β release *ex vivo*. Phenotypic, genomic, and proteomic analysis of the *C. albicans* isolates suggested increased tissue adhesion and invasion of CD-derived strains, which rapidly translocated the epithelial barrier in a dynamic human ‘gut-on-a-chip’ model. Together, these data uncover a novel role for Vδ2+T-cells in restricting the growth of human gut-adapted *C. albicans* strains with virulent features that have been strongly implicated in IBD.

## Results

### Bacterial activation of human intestinal Vδ2+ T-cells restricts growth of fungal pathobionts

Vγ9Vδ2+ (Vδ2+) T-cells are ‘unconventional’ lymphocytes that uniquely respond to BTN3A-mediated sensing of the microbial metabolite HMB-PP^23,24^, which is produced by a wide range of human commensal bacteria but not fungi^23^. We previously reported that Vδ2+ T-cells can promote IL-22 expression by conventional gut T-cells and induce calprotectin release in human mucosal biopsy cultures^25^, suggesting a likely role in anti-microbial barrier defense. To explore the biological relevance, we established human intestinal organ cultures with intact epithelium, which is the primary target of IL-22 signaling^26^. Strikingly, we observed that human mucosal biopsies cultured under standard conditions displayed frequent outgrowth of endogenous fungi, despite extensive washing of the tissue fragments, but not when cultures were supplemented with HMB-PP to selectively activate Vδ2+ T-cells^23^ (**Fig 1A** and **B**). Indeed, HMB-PP activation of gut Vδ2+ T cells was able to restrict growth of a variety of fungal species isolated from human intestinal tissue, including a range of thermotolerant yeasts and moulds (**Fig 1C-F**), the most prevalent of which was *C. albicans* (11 of 17 isolates). HMB-PP had no direct effect on fungal growth when added to cultures in the absence of human cells (data not shown). While other authors have reported that food-associated yeast *Debaryomyces hansenii* colonises regions of inflamed gut in Crohn’s disease (CD) and delays mucosal wound healing in a murine injury model^27^, we did not recover this species using our biopsy culture-based method. These data are consistent with reports that *C. albicans* dominates the subset of specialized fungi that colonize the intestinal mucus layer, whereas a more diverse range of transient species can be detected in the gut lumen^28^.

**Figure 1.**
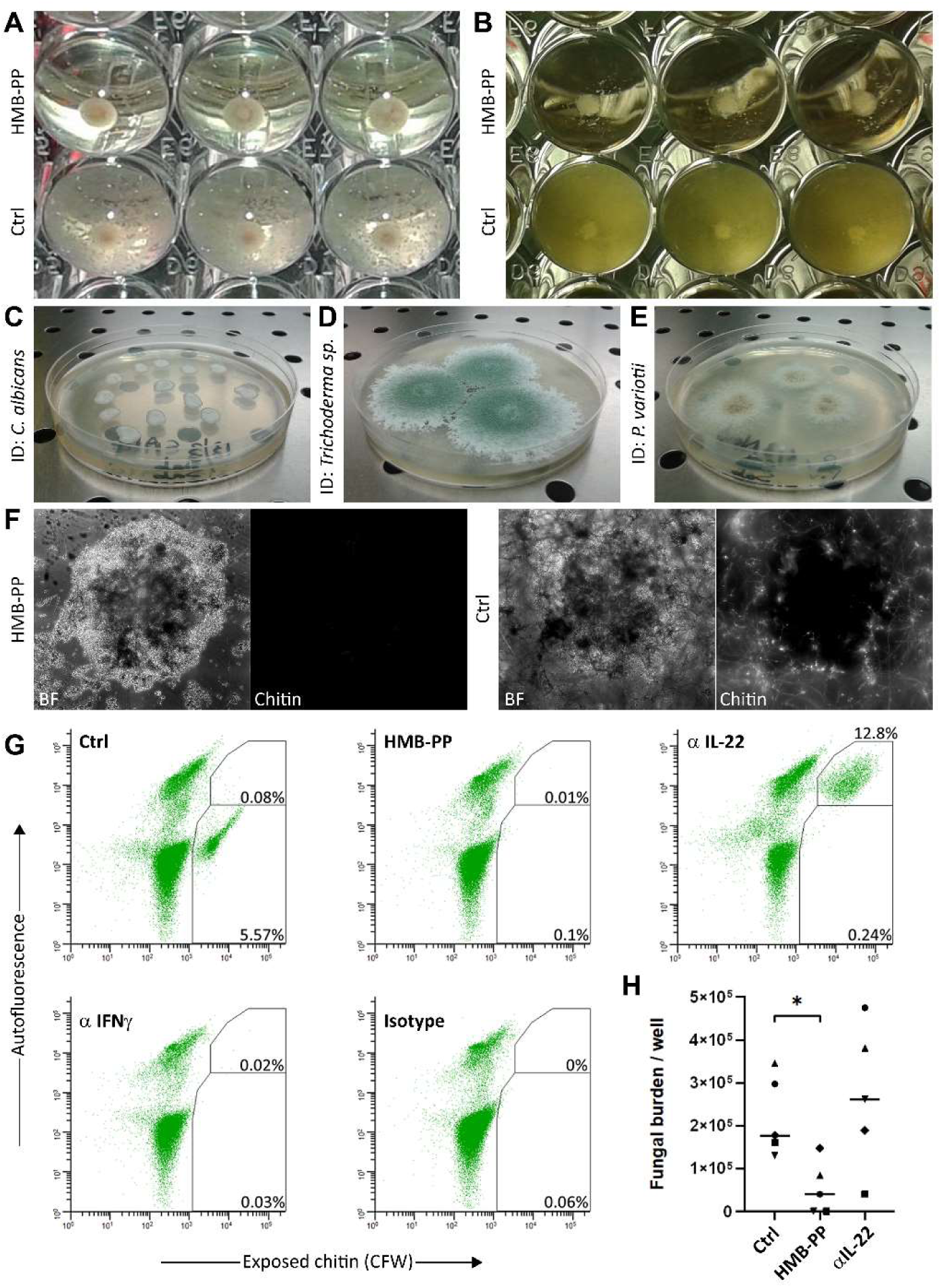
Bacterial activation of human intestinal Vδ2+T-cells suppresses fungal growth. Human intestinal organ cultures were supplemented with epithelial damage-associated cytokine IL-15 and stimulated or not with bacterial phosphoantigen HMB-PP (10nM) for up to 3 days at 37°C, which restricted outgrowth of endogenous fungi (A and B), including polymorphic species *C. albicans* (C) and thermotolerant moulds such as *Trichoderma sp.* (D) and *Paecilomyces variotii* (E). Images show round-bottom chambers of a 96-well plate containing biopsy-derived cells after 3 day culture either photographed from below (A and B), cultured on Sabouraud agar (C, D, E) or stained with calcofluor white prior to wide-field imaging at x20 magnification (F) to visualize cellular structures (brightfield; BF) and exposed chitin fluorescence. Supplementation of cultures with neutralising anti-cytokine antibodies and subsequent analysis by flow-cytometry revealed a critical role for IL-22 in Vδ2+ T cell-mediated fungal control, but not IFNγ or an isotype-matched control antibody (G). Quantification of fungal burden in gut biopsy cultures from n = 5 control donors confirmed that HMB-PP induced significant restriction of endogenous fungal growth (p = 0.0178 by ANOVA), but not when IL-22 was neutralised (H).

We have previously shown that Vδ2+ T-cell activation in human mucosal biopsy tissue upregulates colonic CD4+ T-cell expression of IL-22 and IFNγ, both of which have been implicated in different phases of host antifungal defense^29^. Therefore, we used blocking antibodies to test which of these mediators was critical for fungal suppression in human gut biopsy cultures. While neutralization of IFNγ was indistinguishable from treatment with an isotype-matched control antibody, which had little impact on fungal outgrowth, blockade of IL-22 severely disrupted Vδ2+ T cell-mediated fungal control (**Fig 1G** and **H**). These findings are in-line with previous data showing that IL-22 signaling in the gut epithelium represents an important mechanism of colonization resistance against opportunistic pathogens^30^, as well as our earlier report that Vδ2-induced *AHR* expression and IL-22 signaling are associated with mucosal release of antifungal calprotectin^31,32^. Together, these data reveal that activation of human intestinal Vδ2+ T-cells by bacterial metabolite HMB-PP can suppress fungal pathobionts in healthy gut tissue via upregulation of IL-22.

### Vδ2+ T cell-mediated fungal suppression is impaired in patients with Crohn’s disease

In addition to being a dominant member of the human gut mycobiota, *C. albicans* has been strongly implicated in the pathogenesis of inflammatory bowel disease (IBD)^15^, which is associated with genetic variants that impact major fungal receptors and pathways (including Dectin-1/CLEC7A, NOD2, and CARD9^33–37^) as well as IL-22 signaling in the gut epithelium^38–40^. Indeed, we previously reported that Vδ2+ T-cells can potently upregulate IL-22 in healthy blood and gut lymphocytes^25^, whereas here we observed that Vδ2+ T-cells from IBD patients displayed significantly impaired antigen-presenting phenotype and IL-22 induction in active compared with inactive disease (**Fig 2A** and **B**). These findings are in addition to our previous finding that CD patients receiving azathioprine therapy exhibit very low frequencies of Vδ2+ T-cells in blood and intestinal tissue^41^. Accordingly, unlike biopsy cultures from healthy donors, gut tissue from CD patients contained only trace numbers of Vδ2+ T-cells, which did not expand significantly upon HMB-PP treatment (**Fig 2C**) and failed to limit growth of endogenous fungal species or protect cultures spiked with highly pathogenic *C. albicans* reference strain SC5314 (**Fig 2D** and **E**). These findings are consistent with murine data showing that IL-22 can restrain *C. albicans*-induced release of pro-inflammatory IL-1β^42^, which correlates with colitis severity in human patients^15^, whereas actively inflamed mucosal tissue is instead depleted of IL-22-producing T-cells^43^. Together, these results indicate that impaired Vδ2+ T-cell frequency and function in CD results in failure of bacterial HMB-PP to induce IL-22 expression and restrict intestinal fungal growth.

**Figure 2.**
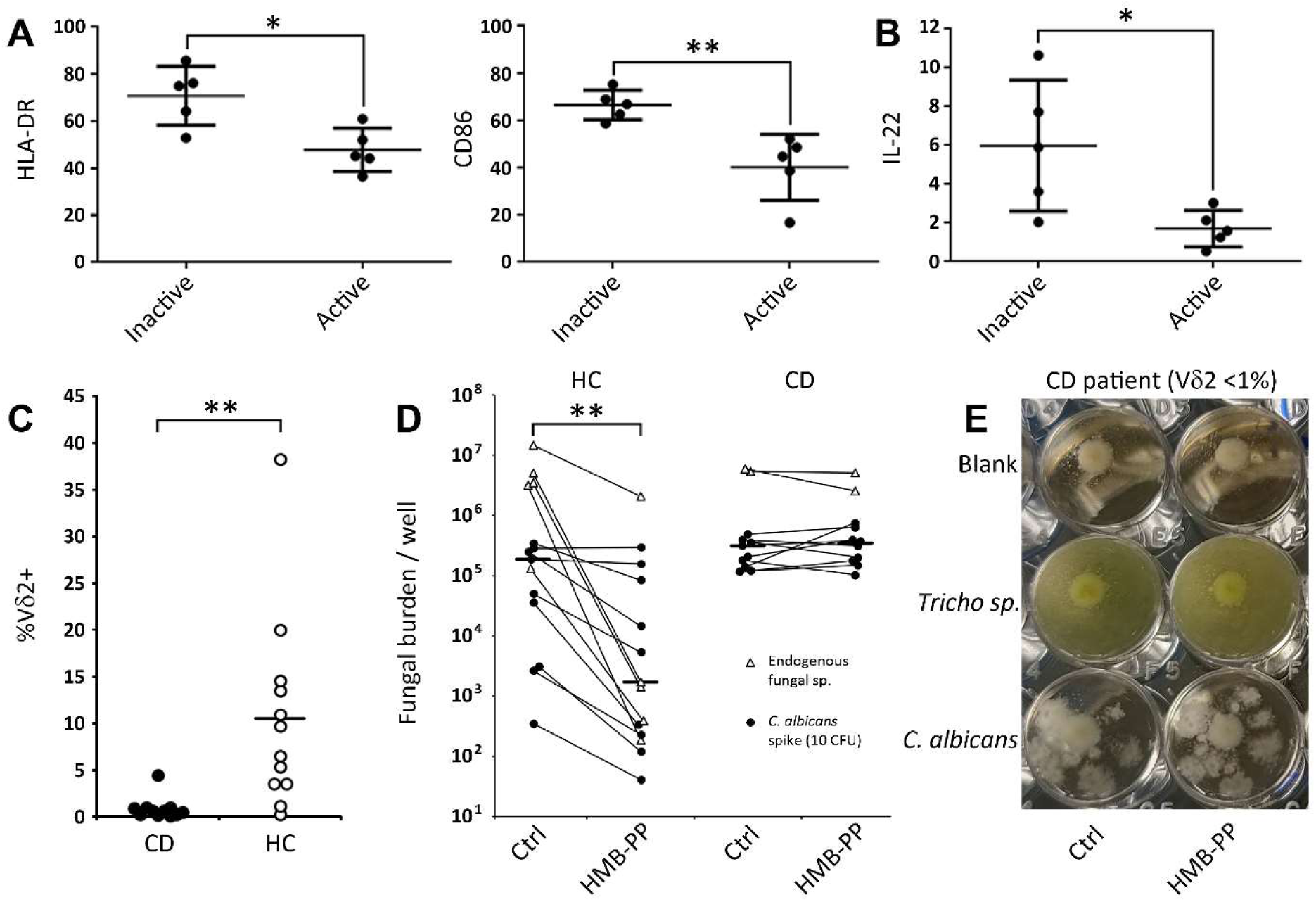
Vδ2+ T cell-mediated fungal suppression is impaired in Crohn’s disease patients. When stimulated with 10nM HMB-PP for 3 days at 37°C, blood Vδ2+ T-cells from IBD patients were significantly less responsive in active compared with inactive disease (n = 5 per group), as indicated by reduced expression of antigen presentation markers HLA-DR and CD86 (A), as well as poor induction of IL-22 in allogeneic naïve CD4+ T-cell responders from healthy donors (B). Gut biopsy tissue from patients with CD also contained low levels of mucosal Vδ2+ T-cells that failed to proliferate during 3 day culture with HMB-PP (C; n = 11, mean 0.85% of T-cell pool), unlike the significant expansions observed in stimulated cultures of healthy gut tissue (mean 10.56%). Accordingly, HMB-PP activation of healthy gut biopsy tissue (n = 13) potently restricted growth of endogenous fungi or spiked-in *C. albicans* reference strain SC5314, but did not reduce fungal burden in cultures of mucosal tissue from CD patients (D and E, n = 11). Bars indicate mean values throughout (with standard error in A and B). *p<0.05, **p<0.01 by T-test (A, B, C) or Wilcoxon Signed Rank Test (D).

### Biodiversity of *C. albicans* strains isolated from healthy and inflamed human intestine

Previous studies have been unable to link changes in mycobiota composition with IBD severity, but increased gut burden of *Candida* has been a consistent finding^44^. We obtained a range of different fungi from human gut biopsy cultures, among which *C. albicans* was the species isolated most frequently (**Fig 3A** and **B**). In a recent analysis of fungal isolates from human mucosal lavage, *C. albicans* strains that readily formed hyphae induced greater levels of mouse macrophage damage *in vitro* and colonic injury *in vivo*, which required the filamentation regulator *EFG1* and CLYS-encoding gene *ECE1*^15^. We therefore assessed filamentation capacity, *ECE1* gene expression levels, and extent of gut epithelial cell damage inflicted by our collection of human gut biopsy-derived strains.

**Figure 3.**
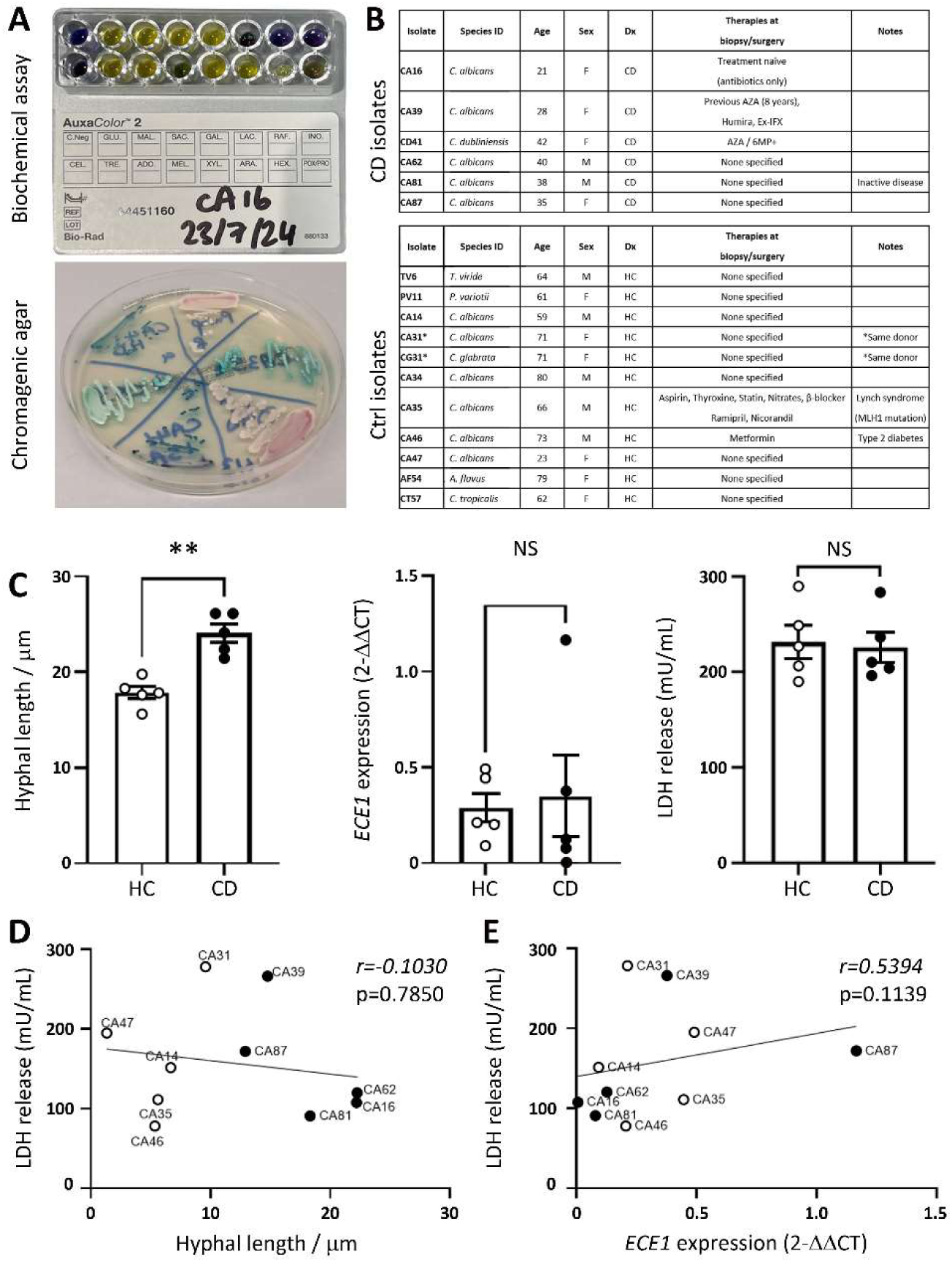
Biodiversity of *C. albicans* strains from healthy and inflamed human intestine. Fungi isolated from human intestinal tissue were identified using a range of morphological and biochemical criteria, including AuxaColor2 biochemical and CHROMagar *Candida* assays (A). *C. albicans* was the most prevalent species in both healthy control and Crohn’s disease (CD) biopsy cultures (B). The *C. albicans* gut strains were next assessed to determine extent of filamentation on plastic, as well as *ECE1* gene expression, and LDH release as a measure of epithelial damage during co-culture with Caco-2 cells (C). CD-derived strains displayed increased hyphal length after 2 h culture (**p = 0.0079 by Mann-Whitney test), whereas qPCR analysis of *ECE1* mRNA indicated comparable expression levels between groups. Fungal co-culture with Caco-2 cells indicated similar damage induction by *C. albicans* strains from HC and CD donors under basic conditions *in vitro*. Further assessment using linear regression did not directly correlate epithelial damage with either germ-tube/hyphal length (D) or *ECE1* mRNA expression level (E). Parts C, D, and E show *C. albicans* strains from donors <75 years-old to limit the impact of advanced age (which in primates impacts known influences on fungal biology including gut pH, motility, oxygen level, and IgA secretion^113^). HC-derived *C. albicans* strains are shown as white circles (n = 5), and strains from CD patients as black circles (n = 5). Bars in part C indicate mean and standard error.

Universal PCR primers (based on 350 *C. albicans* genome sequences) were designed to detect regions of the *ECE1* gene that are conserved across *C. albicans* strains. All human intestinal *C. albicans* strains derived from biopsy cultures expressed detectable *ECE1* mRNA, albeit at highly variable levels that did not reflect patient donor status (**Fig 3C**). Importantly, CLYS production, processing, and delivery via the hyphal tip is a complex multi-step process that cannot be fully appraised at the level of *ECE1* transcription^45,46^, so we proceeded to investigate other key virulence factors including hyphal extension and damage induction. These analyses revealed that CD-derived strains exhibited significantly greater hyphal extension during culture *in vitro*, although this was not significantly correlated with damage inflicted on gut epithelial cells in co-culture assays (**Fig 3C** and **D**). Similarly, *ECE1* mRNA expression levels did not directly predict gut epithelial cell damage *in vitro* (**Fig 3E**). These findings are consistent with reports that *C. albicans* SC5314 is more pathogenic than typical clinical isolates, for which *ECE1* gene expression only partially correlates with epithelial damage *ex vivo*^47^. Together, these data suggest that host microenvironmental factors interact with *C. albicans* genotype to modify patterns of filamentation, hypha maintenance, *ECE1* gene expression, and CLYS export in the human intestine^46^.

### Human gut-adapted morphotypes are shaped by genetic and microenvironmental factors

To better understand the drivers of *C. albicans* biodiversity in healthy and CD intestine, we employed whole genome sequencing of the gut-adapted strains for comparison with the Pasteur reference collection of clinical isolates^48^. When SNP distributions were assessed across our gut strain collection, we observed a wide range of phylogenetic relationships between isolates which spanned multiple different clades (**Supplementary Fig. 1**), suggesting that a wide range of genotypes are compatible with human gut colonization. Given our previous observation that CD-derived strains exhibited greater hyphal extension during culture *in vitro*, we next focused our analysis on regulators of the core filamentation response. In laboratory strains of *C. albicans*, this program is controlled by 8 genes (*ALS3, DCK1, ECE1, HGT2, HWP1, IHD1, RBT1,* and *orf19.2457*)^49^, although mutations in numerous downstream genes can also influence fungal adaptation to the mammalian gut^50–52^. All gut biopsy-derived strains in our collection featured SNPs in one or more regulators of the core filamentation response (**Fig. 4**), irrespective of whether they were isolated from healthy control donors or patients with CD. Indeed, data from other labs indicate that *C. albicans* hyphal genes can be expressed by both yeast and filamentous growth forms during colonization of the mouse gut^50,53^, reflecting that host environment exerts a strong influence on intestinal phenotypes. A key mechanism of *C. albicans* control in the intestine is oxygen availability, which in healthy gut decreases from duodenum to colon, where the epithelium is maintained in a state of physiological hypoxia (<1% O_2_) to restrict pathogen growth^54,55^. To gain further insight into the host factors that shape human gut-adapted morphotypes, we used pSILAC proteomics to probe functional differences between exemplar yeast-like and hyphal strains under ‘gut-like’ hypoxic conditions. For this analysis, we selected the highly filamentous strain CA16 (from a 23 year-old female patient with severe CD who had received only antibiotic treatment) and closely donor-matched isolate CA47 (from a healthy 21 year-old female donor on no medication) (**Fig 5A and B**). Among a list of >2500 fungal proteins identified (**Supplementary Table 1 online**), this analysis revealed starkly different biological profiles for each isolate, with CD-derived strain CA16 being significantly more responsive to environmental change (>900 proteins upregulated in hypoxic culture versus <50 for HC strain CA47; p<0.05 by one-way ANOVA). These data are consistent with reports that commensal organisms are less sensitive than pathogens to varying levels of heat and oxygen, which are typically stable in their natural host environment^56^. Indeed, key processes in hypoxic CA16 included ‘filamentous growth’, ‘negative regulation of immune response’, and ‘sequestering of zinc ion’, which are key determinants of *C. albicans* translocation across intestinal barriers^57^ (**Supplementary Fig. 2**). Strikingly, filamentation was not a highlighted pathway in hypoxic CA47, which was instead characterized by symbiont modulation of host immunity.

**Figure 4.**
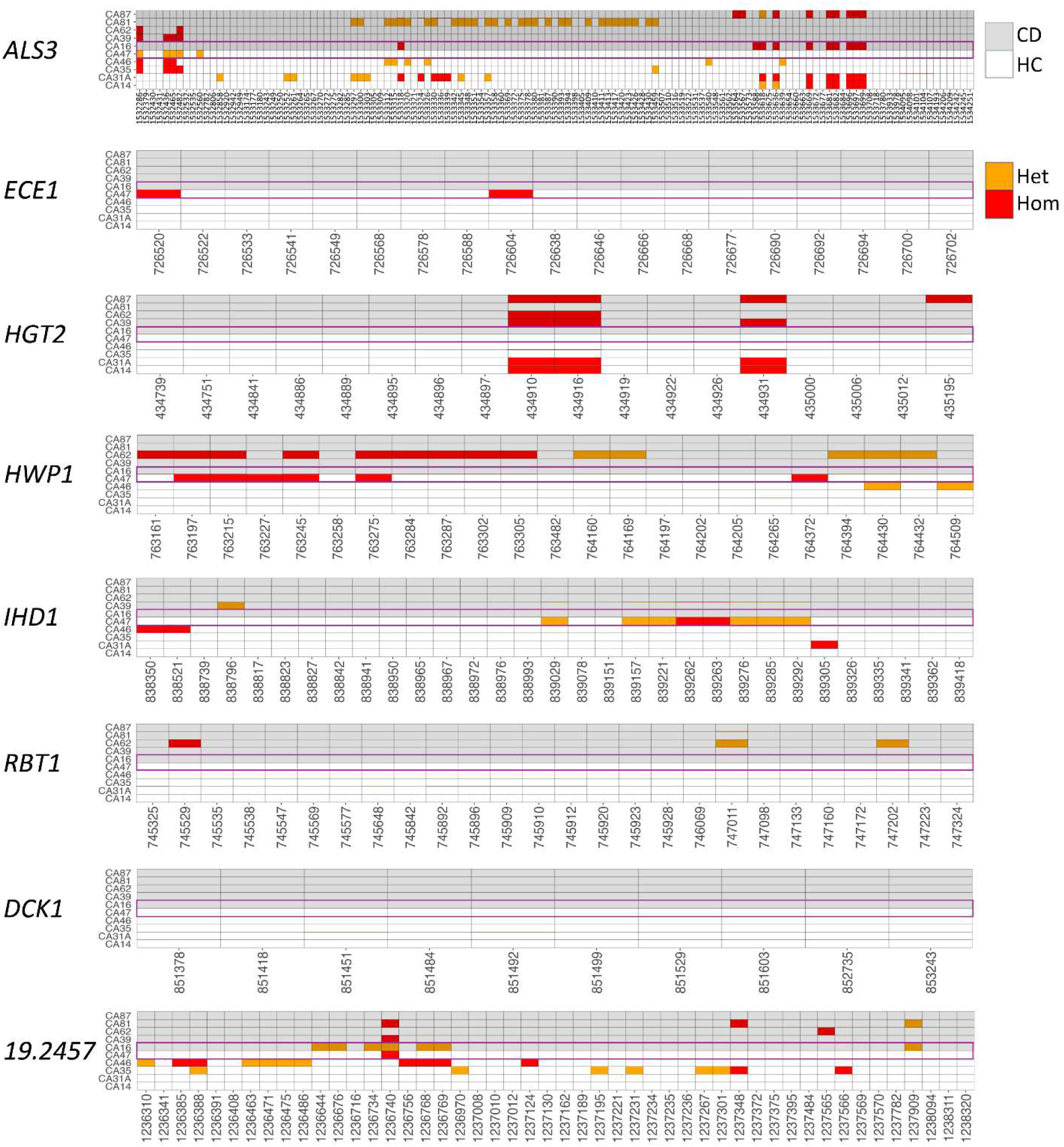
Human gut-adapted *C. albicans* strains exhibit core filamentation gene mutations. Gut-adapted strains were subjected to whole genome sequencing by Illumina methodology and variants were called against the *C. albicans* reference genome (Ca22) using the GATK *HaplotypeCaller* pipeline. All gut biopsy-derived strains featured gene mutations in one or more regulators of the core filamentation response^49^ (Orange = heterozygous, Red = homozygous). Purple boxes highlight SNPs in exemplar strains HC-CA47 and CD-CA16 selected for subsequent functional experiments.

**Figure 5.**
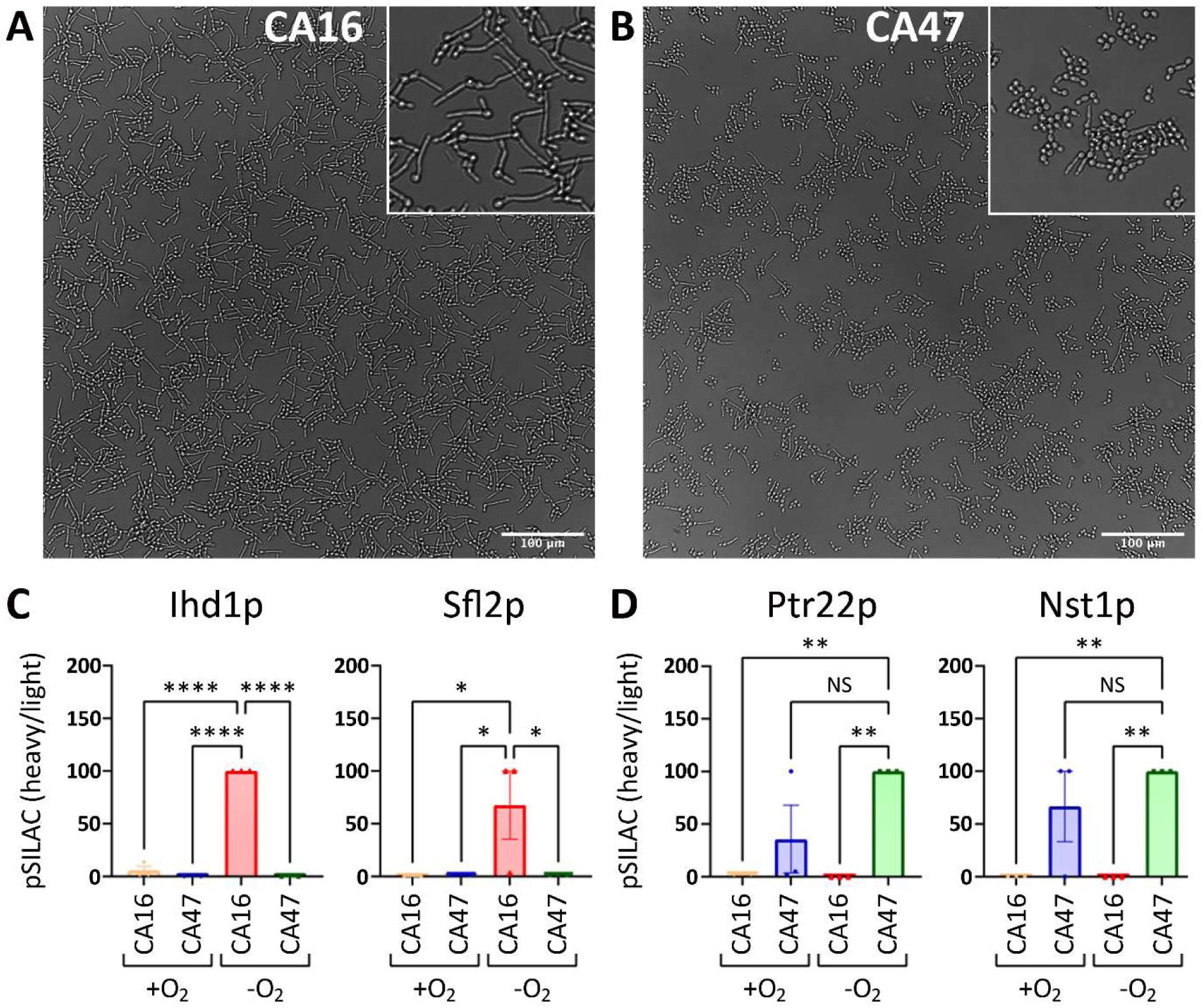
Hypoxia modifies the biology of gut-adapted yeast and filamentous *C. albicans*. Representative CD-derived strain CA16 (A) and HC-derived strain CA47 (B) were cultured in YNB medium (supplemented with sucrose and ammonium sulphate) in either normoxia or hypoxia (∼98% N_2_, ∼2% O_2_) overnight, then changed to pSILAC medium (R/arginine and K/lysine-free RPMI medium supplemented with heavy R6 and K8 amino acids) for a further 3 h prior to analysis by pSILAC proteomics. In low-oxygen cultures resembling human gut conditions, the top 10 differentially expressed proteins included hyphal regulators Ihd1p and Sfl2p, which were actively translated only by CD-derived strain CA16 (C). HC yeast CA47 instead expressed mediators of niche-specific stress tolerance, including Ptr22p and Nst1p (D). Scale bars indicate 100μM (A and B). Bars in C and D indicate mean and standard error of n = 3 biological repeats (*p<0.05, **p<0.01, ****p<0.0001 by ANOVA).

Together, these data suggested that CD-derived strain CA16 is poised for filamentation and pathogenesis in conditions that resemble the human gut environment. Among the proteins most strongly expressed by CA16 during hypoxic culture was core filamentation regulator Ihd1p (**Fig 5C**), whereas this hyphal-specific protein was barely detected in CA47 (consistent with their contrasting SNP profiles for this gene; **Fig 4**). A similar pattern was observed for hyphal regulator Sfl2p which senses microenvironmental factors to promote virulence in mouse gastrointestinal infection^58,59^, and was strongly expressed by CA16 but not CA47 despite both strains lacking SNPs in the corresponding gene. Conversely, CA47 was enriched in regulators of nutrient uptake and tolerance of tissue-specific variables including iron, oxygen, heat, and osmotic stress^60^ (**Fig 5D**), as well as epigenetic adaptation to the mammalian gut e.g. Med9p^50,61^ (**Supplementary Table 1 online**). These data resemble reports that colonization of the mouse oral cavity is associated with attenuation of the hyphal programme and metabolic adaptation to local conditions, although capacity to filament at the host-fungal interface remains intact^62^. Strain-dependent pathogenicity is therefore shaped by a complex interplay of fungal genetics and niche-specific factors that likely include variable host immune responses.

### Phagocyte responses to intestinal *C. albicans* are modified in inflammatory bowel disease

Neutrophils and monocyte-macrophages are critical mediators of antifungal immunity and dominate the inflammatory infiltrate in UC and CD respectively^63,64^. Key functions of these lineages include neutrophil engulfment of fungi and hyphal destruction via NETosis^65^, alongside macrophage phagocytosis and folding of developing hyphae^66^. Intestinal phagocytes also play a key role in inducing mucosal IgA responses to fungal hyphae^67,68^, with the IgA2 subclass promoting NET formation and macrophage cytokine release^69^, while patients with selective deficiency in this subclass exhibit increased risk of ileal CD^70^. Colitis severity has also been linked with extent of *C. albicans*-induced murine macrophage injury and release of IL-1β^15^. We therefore assessed the induction of NETosis by our human gut-derived *C. albicans* strains, as well as monocyte-derived macrophage (MoMΦ) susceptibility to CLYS exposure, strain-dependent damage, and IL-1β induction *in vitro*.

When co-incubated with healthy donor blood neutrophils, CD-derived strain CA16 triggered significantly higher levels of NETosis relative to HC-derived strain CA47 (**Fig 6A**), consistent with the known ability of neutrophils to discriminate between yeast and hyphal morphologies^65^. Intriguingly, divergent NETosis responses to CA16 and CA47 were lost in assays using blood neutrophils from CD patients (**Fig. 6B**), suggesting immaturity and/or functional impairment of this compartment in disease^71,72^. In-line with these findings, chemotherapy-induced neutropenia or functional defects in this compartment are known to confer increased risk of fungal infections e.g. chronic granulomatous disease (CGD)^73^. Similarly, humans with *STAT1* gain-of-function mutations exhibit chronic mucocutaneous candidiasis (CMC), which is associated with reduced numbers of classical monocytes that display altered patterns of microbial sensing^74^. When we performed similar assays using healthy blood MoMΦ, these cells released only limited amounts of IL-1β upon challenge with HC strain CA47, which was indistinguishable from exposure to a CLYS toxin-only control (which lacks potency when not delivered to host cell membranes via hyphal invasion pockets^47^) (**Fig 6C**). In contrast, CD-derived strain CA16 was a strong inducer of MoMΦ-derived IL-1β, which was released at comparable levels to macrophages challenged with hyper-virulent reference strain SC5314 (**Fig 6D**). Similar patterns of IL-1β release were observed when using MoMΦ generated from patients with CD, although the efficiency of fungal clearance was markedly reduced, irrespective of the strain used in co-culture assays. Together, these data indicate that key human leukocyte populations mediating antifungal immunity and mucosal inflammation display altered responses to *C. albicans* strains derived from CD intestine compared with healthy gut tissue.

**Figure 6.**
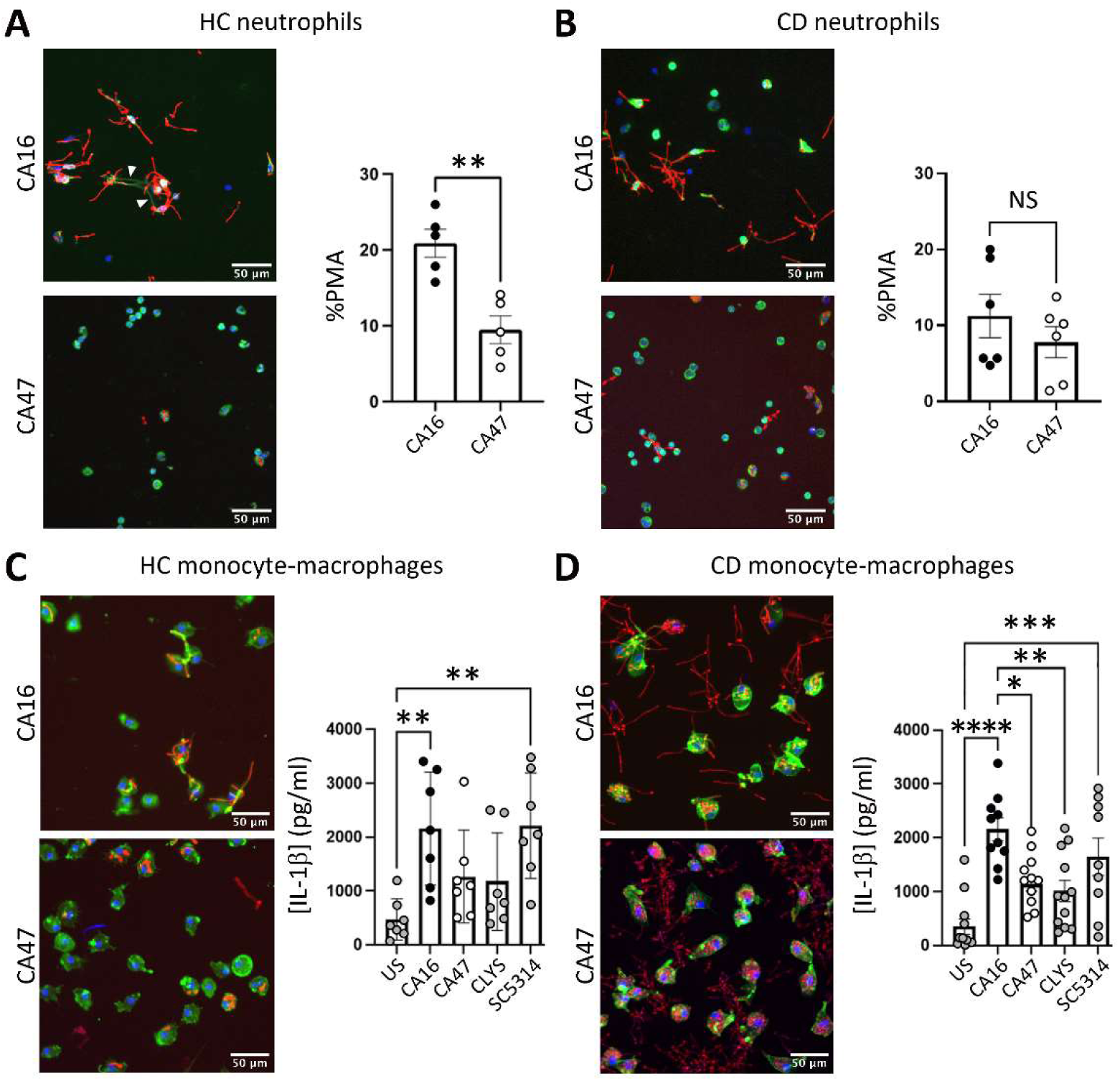
Phagocyte responses to human intestinal *C. albicans* strains are modified in CD. Peripheral blood from HC and CD donors was used for isolation of CD66b+CD16+ neutrophils or CD14+ classical monocytes that were differentiated into macrophages by 7 day culture with M-CSF. Neutrophils were incubated with fluorescent *C. albicans* isolates (red) for 3 h before staining with antibodies against MPO (green), and calprotectin (grey), with Hoechst to visualise nuclei (blue). Production of neutrophil extracellular trap (NET)-like structures was quantified relative to the PMA positive control using SYTOX Green assays. (A) Neutrophils from HC donors (n = 5) produced greater numbers of NET-like structures (white arrows) when challenged with CA16 rather than CA47 (p = 0.0079; unpaired t-test and Mann Whitney test). (B) In contrast, neutrophils from IBD patients (n = 6) produced similar levels of NET-like structures when exposed to either CA16 or CA47 (p = 0.4848; unpaired t-test and Mann Whitney test). Data are presented as mean ± SEM. Similarly, blood monocyte-derived macrophages were primed with 50 ng/mL LPS from *Klebsiella pneumoniae* for 2 h then challenged with CLYS synthetic toxin or CA16, CA47, or SC5314 (all MOI 5) for 5 h before assessing IL-1β release by ELISA. Representative images show fluorescent *C. albicans* strains (red), Hoechst-stained nuclei (blue), and phalloidin-stained actin filaments (green). When co-cultured with macrophages from healthy donors (n = 7), CD strain CA16 induced comparable levels of injury/IL-1β release to reference strain SC5314, whereas HC strain CA47 was indistinguishable from the CLYS-only control (C). While similar patterns were obtained using macrophages derived from CD donors (n = 10), fungal burden per well remained higher in these assays (D). Scale bars indicate 50μM. Graph bars indicate mean and standard error. *p<0.05, **p<0.01, ***p<0.001, ****p<0.0001 by T-test (A and B) or ANOVA (C and D).

### Rapid fungal adhesion and invasion of the gut barrier in a human ‘organ-on-a-chip’ model

Typical mouse models of fungal colonization rely on antibiotic treatment to deplete gut bacteria, which allows yeast morphotypes to out-compete hyphal cells^52,67^. In this setting, the strain-dependent kinetics of epithelial binding and invasion depth exert a critical influence on *C. albicans* commensalism versus tissue damage and expulsion from the gut^47^. However, recent data suggest that strains able to undergo hyphal transition can be significantly fitter than yeast-locked cells when the bacterial microbiota is intact^75^, suggesting a major influence of gut environmental factors on fungal colonization. Our proteomic data also suggested variable tissue adhesion/invasion potential of strains CA16 and CA47 under ‘gut-like’ conditions *in vitro*, but it was unclear whether this might apply under dynamic conditions in the human intestine *in vivo*. Since *C. albicans* does not naturally colonize mouse intestine, we instead developed a human ‘gut-on-a-chip’ model to explore fungal adhesion/invasion dynamics in the relevant ‘*in vivo*’ setting. We first transformed exemplar clinical strains CA16 and CA47 with pENO1-iRFP-NATr fluorescent reporters (as previously described^76^) then assessed fungal binding and translocation across the model gut barrier when subject to physiologically relevant forces of liquid flow and peristaltic stretch (**Fig 7A-E**).

**Figure 7.**
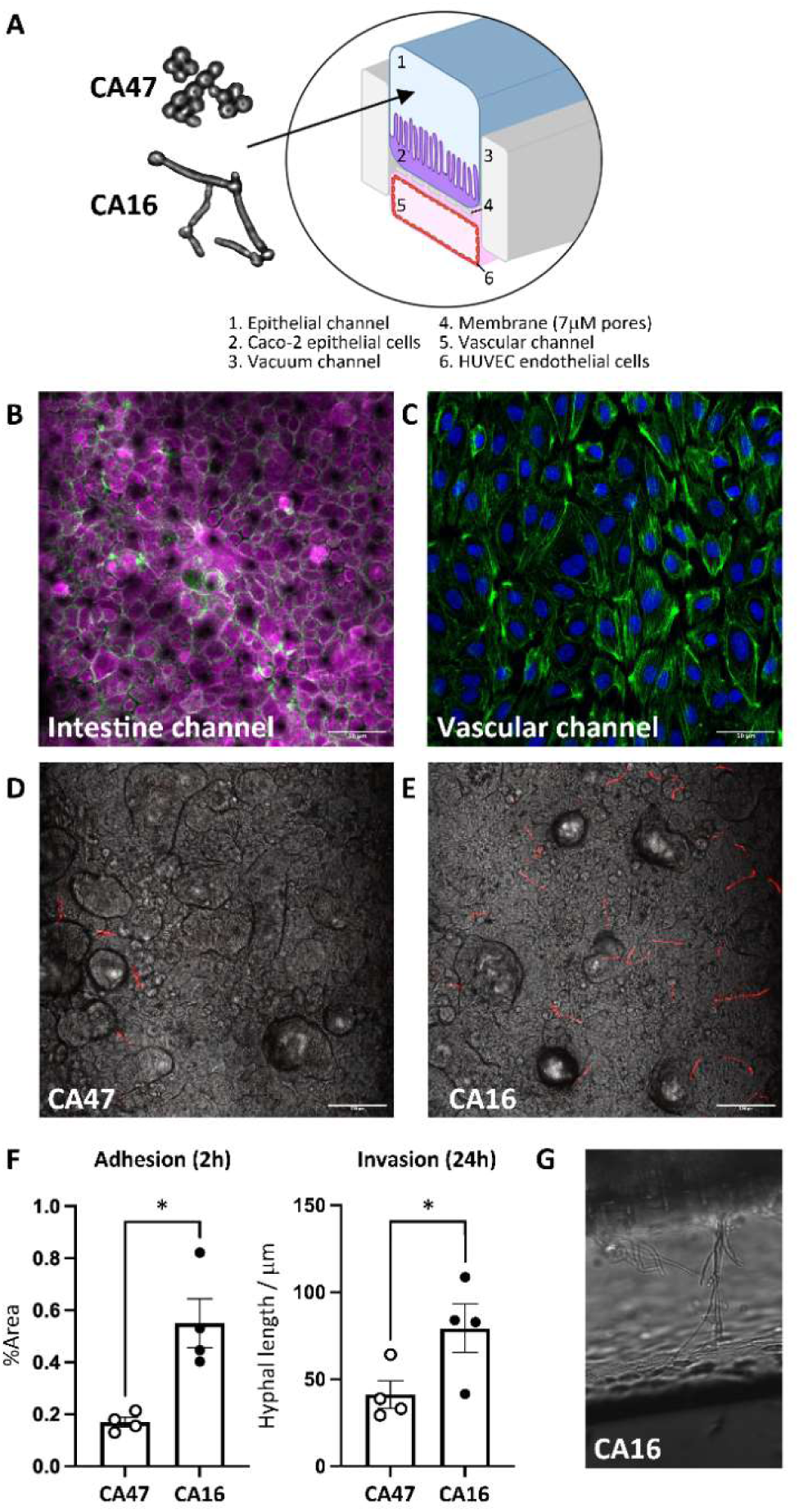
Rapid fungal adhesion and invasion of a dynamic human ‘gut-on-a-chip’ model. Schematic of the gut-on-a-chip model formed of two parallel channels separated by a porous membrane and subjected to physiologically-relevant fluid flow and stretch forces (A). The intestinal channel contained Caco-2 epithelial cells which formed continuous tight junctions and 3D villous-like structures under peristaltic motion (B). The underlying vascular channel was coated with HUVEC endothelial cells to create a model blood vessel (C). Human gut-adapted *C. albicans* strains with pENO1-iRFP-NATr fluorescent reporters were then introduced to the model gut lumen for assessment of host tissue binding and invasion potential. Early-stage adhesion was calculated as % surface area bound by fluorescent *C. albicans* after 2 h incubation using microscopy and Fiji software, which revealed limited binding of HC yeast CA47 (D) relative to filamentous CD strain CA16 (E), which was significantly more adhesive by unpaired t-test (F; p = 0.0074, n = 4 per group). Indeed, when hyphal protrusions into vascular channel Z-stack images were measured at 24 h post-infection, CD strain CA16 was also significantly more invasive (p = 0.0412, n = 4 per group), and displayed extensive growth into the model blood vessel (G; magnification x20). Scale bars indicate 50μM (B and C) or 100μM (D and E). Bars in F indicate mean and standard error (*p<0.05).

Tissue 3D structure and peristaltic motion have been shown to modify invasion of the primate-specific bacterial pathogen *Shigella flexneri* into human gut models^77^, and key genes associated with fungal adhesion to biological surfaces can be expressed differentially between strains as early as 90 min after contact^78^. We therefore assessed epithelial binding 2 h after spiking human gut-adapted strains CA16 and CA47 into our intestine-on-a-chip system under conditions of physiological flow and mechanical stretch (**Supplementary Video 1**). When quantified by fluorescent imaging, commensal-like HC strain CA47 displayed only low levels of adhesion to the gut surface after 2 h incubation, with limited filamentation *in situ*. In contrast, CD-derived strain CA16 exhibited rapid binding to the gut surface with extensive hyphal growth, followed by significant translocation into the underlying vascular channel within just 24 h (**Fig 7F, G, and Supplementary Video 2**). These functional assays were consistent with our genotypic, proteomic, and immunological data suggesting that CD-derived strain CA16 is more pathogenic than healthy control-derived strain CA47. Importantly, both strains were capable of binding to the gut epithelium and eventually invaded the model vasculature even when subjected to dynamic physiological forces. Future work will therefore require the development of more advanced organ chip models, replete with multiple lineages of human leukocytes, which can be used to determine how these fungal dynamics are modified by host immunity in health and inflammatory settings ‘*in vivo*’.

## Discussion

Here we report that bacterial metabolite HMB-PP can activate human intestinal Vδ2+T-cells to suppress fungal pathobionts in healthy gut tissue via an IL-22-dependent mechanism that is dysregulated in Crohn’s disease (CD). These data are in-line with previous reports that commensal metabolites can modify host immunity to influence competition between microbial strains, species, and even kingdoms^3,5–7^, whereas defects in BTN/L regulation of the γδ T-cell compartment have been linked with severe CD phenotypes^79^. Indeed, previous studies in animals have shown that hundreds of blood biochemicals are present only after microbe colonization^1,2^, and may influence host physiology even more strongly than heritable factors^2,80^. However, the specific microbial species involved, immune mechanisms they trigger, and consequences for human body tissues have remained unclear.

γδ T-cells play a critical role in epithelial defense against microbial invasion in the mouse intestine^81^, where IL-22 is known to play a dominant role in the early response to adherent-invasive pathogens^29,82^. While T-cells producing IL-17 are strikingly increased in inflamed human gut, those producing IFNγ and IL-22 are significantly depleted, in correlation with relative loss of the bacterial phylum Firmicutes^43^ (which includes numerous HMB-PP-producing genera^23^). These data are consistent with our previous finding that bacterial activation of Vδ2+T-cells in human gut tissue does not alter IL-17 expression but instead induces *AHR* and IL-22^25^, which in the current report emerges as a potent mechanism of fungal suppression in human gut biopsy cultures. Indeed, other authors have also observed a key role for anaerobic Firmicutes in preventing *C. albicans* colonization of mouse gut via activation of hypoxia-inducible factor (HIF)-1α, which cross-talks with the AhR pathway to promote T-cell expression of IL-22 but not IL-17^83–85^. Our results further suggest that this axis is disrupted in CD, since patient blood and biopsy tissue were characterised by poor Vδ2+T-cell sensitivity to HMB-PP and trace cell numbers in intestinal mucosa, accompanied by rapid outgrowth of endogenous fungi from biopsy cultures. These findings resemble reports that the AhR/HIF-1α/IL-22 pathway is impaired in gut biopsies from IBD patients^86^, whereas provision of AhR ligands or drugs that restore epithelial hypoxia can limit *C. albicans* growth^55^, likely by inducing IL-22 and AMP release^87,88^. Indeed, without a robust Vδ2+T-cell response, our human gut biopsy cultures yielded a range of viable yeasts and moulds that have been reported to cause serious infections in humans, with *Candida* being the most common genus and *C. albicans* the most dominant species. While strains derived from CD gut tissue were significantly more filamentous, *ECE1* gene expression and epithelial damage levels could not be distinguished from control isolates under basic culture conditions *in vitro*. These findings suggest that strain-dependent pathogenicity is influenced by additional factors in the human gut environment that are not easily replicated *ex vivo*.

Host defence against *C. albicans* depends not only on control of inflammatory responses but also microenvironmental factors, with optimal balance of Th1/Th17 immunity appearing to vary by epithelial structure, body site, and prevailing local conditions^89,90^. In human intestine, *C. albicans*-responsive cells are strikingly enriched within the Th17 memory pool and expand during mucosal inflammation, which in CD patients is characterised by the further generation of Th1 cells that cross-react with other fungal species^91,92^. However, the ability of unconventional lymphocytes to direct these responses was previously unknown, including the key mediators involved and downstream impact on antifungal defence. Indeed, while γδ T cell-derived IL-22 plays a dominant role in a mouse oral infection model^89^, γδ T cell-expressed IL-17A appears critical in a hypoxic skin injury model^93^. Similarly, human inborn errors of Th17 immunity are strongly linked with mucocutaneous candidiasis, but patients are typically also deficient in IL-22 and IFNγ, with pathology varying significantly between body sites^94,95^. Antifungal cell types and the mechanisms they engage may therefore differ between host species and local tissue architecture^89,96,97^. Conversely, the biological profiles that confer greatest *C. albicans* fitness are influenced by conditions at different anatomical locations, which may subsequently require tissue-specific defence mechanisms. In-line with this, *C. albicans* colonization of mouse cecum is accompanied by high expression of hyphal-linked gene *ECE1*, despite yeast cell morphology being the dominant growth form in this setting^53^. *C. albicans* lacking *ECE1* also displays poor gut colonization that can be reversed by restoring WT *ECE1* gene into the mutant strain, suggesting that CLYS may favour commensalism in the appropriate host setting^75^. Our data further support this concept that a wide range of *C. albicans* phenotypes are compatible with mammalian gut colonisation^98^, since mutations in core filamentation genes were frequent but diverse in nature, and yielded both yeast and hyphal morphologies that tended to align with donor inflammatory status.

While *C. albicans* colonises the intestine of essentially all healthy adult humans, fungal burden varies significantly between individuals^11^, likely under the influence of mucosal immunity and biochemical factors. Indeed, mice deficient in IL-10 display reduced gut colonisation by *C. albicans*^99^, whereas mice lacking IL-12 are colonized at higher levels^90^. Similarly, in healthy gut the epithelium is maintained in a state of physiological hypoxia (<1% O_2_), but barrier disruption can increase oxygen availability and promote dysbiosis^54^. This likely contributes to increased fungal burden and pathogenicity in IBD^17^, as well as modifying the diversity and abundance of competing bacteria, thus altering the fitness benefits of adopting yeast rather than hyphal morphology and toxin production^75^. Accordingly, our proteomic analysis revealed starkly different profiles for CD strain CA16 compared with HC strain CA47 when cultured under ‘gut-like’ hypoxic conditions. Key points of divergence included regulators of filamentation such as IHD1, and reactivity to tissue-specific variables including iron, oxygen, heat, and osmotic stress. Consistent with these data, mice fed a conventional diet are generally refractory to gut colonisation with virulent reference strains such as SC5314 that readily form hyphae, but persistence can be impacted by multiple factors including sugar intake, antibiotic treatment, and mucosal inflammation^75,100^. Indeed, antibiotic exposure also increases risk of new onset CD in humans^101^, and hyphal strains of *C. albicans* are more prevalent in mucosal washing from affected patients^68^, but until now it has remained unclear how these different morphotypes interact with human gut tissue and resident leukocytes.

Earlier reports identified that genetic defects in immune detection of fungi are associated with severe forms of colitis^33^, whereas antifungal therapy accelerates colonic healing^102^. Many IBD risk genes are now recognised to play important roles in host-fungal interactions^103^. In particular, *CARD9* protects mice against colitis by promoting expression of IL-22^104^, which can be triggered by *C. albicans* proximity to the intestinal epithelium^28^. Similarly, *NOD2* expression is largely restricted to the monocyte-macrophage lineage^105^, which plays a crucial role in fungal sensing at the gut barrier^27,68,106–108^. Indeed, human gut-adapted *C. albicans* strains that inflict high levels of murine macrophage damage *in vitro* are associated with increased colitis severity in the patients from whom they were isolated^15^. Similarly, we observed greater levels of neutrophil NETosis and macrophage IL-1β release in response to *C. albicans* derived from CD gut tissue rather than HC biopsy cultures, reflecting enhanced filamentation and potential for host cell damage in disease. A key mechanism of host defence against fungal invasion is thought to require *CARD9*^+^ macrophages that induce antifungal IgG^108^ and secretory IgA which preferentially targets hyphal cells^68^. In healthy intestine, this disadvantages virulent morphotypes and favours commensal yeast that have mutated hyphal genes^50,52^, whereas sIgA selectivity for hyphal targets is impaired in CD^68^, perhaps contributing to the increased filament length we observed among CD biopsy-derived strains. This may also explain why highly pathogenic reference strains such as SC5314 are typically refractory to colonisation of healthy murine intestine, whereas mucosal strains with filamentation defects such as 529L appear more resistant to host antimicrobial factors^88^. However, it is important to note that local environment nonetheless exerts a major influence on colonisation dynamics, since strain 529L continues to produce hyphae adjacent to the colonic epithelium *in vivo*^88^. We therefore developed a custom ‘gut-on-a-chip’ model to assess the extent to which *C. albicans* isolates from HC and CD gut can bind and damage human host tissues under relevant physiological conditions of peristaltic fluid flow and stretch ‘*in vivo*’. Consistent with our morphometric, genomic, and proteomic data, *C. albicans* isolated from CD tissue displayed significantly greater ability to adhere and invade the model tissues.

It is increasingly clear that a complex range of variables can impact host-fungal interactions in the human intestine, including quantity and diversity of bacterial competition, range of nutrient sources available, and local oxygen concentration. Considering major differences between mouse and primate gut physiology, resident microbiota, and immune profiles^4,110^, it is perhaps not surprising that *C. albicans* can represent a keystone commensal species in humans but not in rodents^4,109^. By extension, natural human hosts may engage mechanisms of fungal control that are not present in rodent systems, such as our current finding that mucosal Vδ2+ T cells can suppress growth of intestinal *C. albicans*. Given that both the target microbe and host immune mediator of this axis are absent in rodents, alternative model systems will be needed to fully understand this biology and perhaps uncover additional new aspects of human tissue defence against fungal infection.

## Methods

### Study participants

Peripheral blood and intestinal biopsies were obtained from patients undergoing colonoscopy as part of routine clinical care to assess CD activity, or for colorectal cancer screening or investigation of rectal bleeding but with no significant endoscopic abnormalities (controls). Additional samples of mucosal tissue were obtained from patients undergoing surgical resection for colorectal cancer, non-inflammatory intestinal motility disorders, or severe CD. Collection of relevant clinical data and biological specimens was performed under the NIHR Research Tissue Bank approval ‘BIORESOURCE STUDIES OF THE DIGESTIVE SYSTEM IN CHILDREN AND ADULTS’ (REC Number: 15/LO/2127). All volunteers gave written informed consent prior to inclusion.

### Peripheral blood cells

Human whole blood was collected into sodium-heparin vacutainers (BD Bioscience) and used for PBMC isolation (below) or directly labelled with monoclonal antibodies for 15 min at room temperature prior to lysing red blood cells by addition of Optilyse C (Beckman Coulter, Bucks). Labelled cell suspensions were washed twice in cold FACS buffer (PBS containing 2% fetal calf serum, 0.02% sodium azide, 1 mM EDTA) and fixed in 1% paraformaldehyde prior to analysis by flow-cytometry.

### PBMC isolation

Peripheral blood was collected into sodium-heparin vacutainers, diluted 2:1 with room-temperature RPMI, and the PBMC fraction was isolated by density gradient centrifugation over Ficoll-Plaque PLUS (GE Healthcare). Mononuclear cells were aspirated from the interface and viability was determined by Trypan Blue dye exclusion (Gibco).

### Neutrophil isolation

Blood was collected from healthy volunteers and IBD patients into K-EDTA tubes and processed using EasySep^TM^ Direct Human Neutrophil Isolation Kits (StemCell Technologies, Inc.) according to the manufacturer’s protocol. A total of 75 x 10^3^ neutrophils in 50 µL phenol-red free RPMI-1640 medium (supplemented with 1% HEPES and 1% FBS) were added into each chamber of a 96-well flat-bottom plate and allowed to settle for 30 min in a 37°C, 5% CO_2_ incubator. Next, 50 µL of pre-diluted stimuli were added to replicate wells of neutrophils, followed by incubation for 3 h in a 37°C, 5% CO_2_ incubator (stimuli included 10^6^/mL *Candida*, 200 nM PMA, 5-75 µM CLYS toxin, 5 µM Ionomycin, 1 µg/mL LPS). After incubation, supernatants were removed and cells were either stained with SYTOX Green or fixed immediately by addition of 4% PFA overnight. The following day, supernatant was discarded, cells were washed 3 times with PBS, and a final volume of 200 µL PBS was added for sample storage until analysis.

### Monocyte-derived macrophages

Positive selection of the CD14+ population from PBMCs was carried out using the EasySep^TM^ Human CD14 Positive Selection Kit II (StemCell Technologies, Inc.) according to the manufacturer’s protocol. Primary blood monocytes were adjusted to a final concentration of 3 x 10^5^ per mL in complete medium (Dutch-modified RPMI-1640 supplemented with 10% FBS, 1% Pen/Strep, 1% L-glutamine, 50 ng/mL M-CSF) and then cultured in 96-well plates at 37°C, 5% CO_2_ for 7 days (half the culture volume was replaced on day 3 with fresh medium containing 50 ng/mL M-CSF). Macrophages were harvested by adding medium containing 1 mM EDTA and placing on ice for 10 min.

### Lamina propria mononuclear cells

Intestinal biopsies were washed in 1 mmol/L dithiothreitol (Sigma-Aldrich) and cultured in 24 well plates in complete medium (Dutch-modified RPMI-1640 medium, 10% FCS, 2 mM L-glutamine, 100 u/mL penicillin, 100 μg/mL streptomycin, 25 µg/mL gentamicin, recombinant human IL-2 [30 u/mL] and IL-15 [20 ng/ml]) in the presence or absence of 10nM bacterial phosphoantigen HMB-PP (Echelon Biosciences) and blocking antibodies (ultra-LEAF purified anti-IFN-γ [clone B27] from BioLegend, or anti-IL22 [22URTI] from eBioscience) for up to 3 days at 37°C, 5% CO_2_. After 3 days, intact tissues were discarded and the egressed leukocytes were labelled with mAb and/or fungal content was quantified by calcofluor white staining (25 μg/mL for 15 minutes) prior to analysis by flow cytometry. Alternatively, the cultures were resuspended into 96-well round-bottom plates for subsequent functional assays.

### Neutrophil SYTOX™ Green assay

SYTOX™ Green was used to quantify cells undergoing NET formation. After 3 h incubation with the indicated stimuli, SYTOX™ Green was added at a concentration of 1 µM and fluorescence was measured immediately using a CLARIOStar plate reader at excitation 500 nm, emission 528 nm. Each stimulus was repeated in triplicate and average values were recorded as mean fluorescence. Total % NET release was calculated by subtracting background fluorescence signal (medium containing SYTOX™ Green only) from the values obtained, and then expressed relative to PMA positive control (determined as 100% NET release).

### Neutrophil immunofluorescent staining

Neutrophils were permeabilized with a solution of 0.1% sodium citrate and 0.1% Triton X-100 in PBS for 10 min at 4°C. After washing 3x with PBS, cells were blocked with 3% BSA and 10% serum (either donkey or goat depending on secondary antibodies used) and incubated for 60 min at 37°C. After washing 3x with PBS, diluted primary antibodies (1:1000 H3Cit, 1:100 NE, 1:250 MPO, 1:200 Calprotectin) were added and cells incubated overnight at 4°C. Cells were washed 3x with PBS before secondary antibodies were added (1:500 dilution) and incubated at room temperature in the dark for 2 h. Cells were washed 3x with PBS. Finally, Hoechst (1:10,000 dilution) was added and incubated for 10 min at room temperature. Cells were washed 3 times and resuspended in PBS for analysis.

### Macrophage infection assay

On completion of the differentiation culture, macrophages were primed for 2 h with 50 ng/mL LPS in serum-free medium before challenge with different *C. albicans* strains (MOI 5) or CLYS toxin prepared in sterile water (75 µM final concentration). After 5 h incubation the exhausted culture medium was collected and cytokine concentration determined using Human IL-1β ELISA MAX kits (Biolegend) according to the manufacturer’s instructions.

### Macrophage immunofluorescent staining

Macrophages were differentiated as described above before adding fluorescent reporter strains of *C. albicans* at MOI 1. After 3 h, plates were fixed with 4% PFA for 15 min at room temperature. Macrophages were permeabilized with a solution of 0.1% sodium citrate and 0.1% Triton X-100 in PBS for 10 min at 4°C. After washing 3x with PBS, diluted AF488-Phallodin (1:500) was added and the cells incubated for 2 h at room temperature. Cells were then washed 3x with PBS before adding Hoechst (1:10,000 dilution) and incubating for 10 min at room temperature. Cells were washed 3 times and resuspended in PBS for analysis.

### γδ T-APC assays

Vγ9^+^ and Vδ2^+^ T cells (each >99% purity) were isolated from PBMC using anti-Vγ9-PECy5 (Beckman Coulter) or anti-Vδ2-PE mAb (BD Biosciences), combined with anti-PE microbeads (Miltenyi). γδ T-APCs were generated by co-culture with irradiated monocytes (50 Gy) at a 10:1 ratio in the presence of 10 nM HMB-PP with or without 20 ng/ml IL-15. γδ T-APCs were cultured for 3 d and purified either by positive selection or cell sorting (purity >99%). Bulk CD4^+^ T cells (>95% purity) were isolated from PBMC via negative selection using the CD4^+^ T cell Isolation Kit (Miltenyi) then labeled with anti-CD4, anti-CD45RA, and anti-CCR7 mAb prior to sorting on a FACSAria II (BD Biosciences) to purify the naïve population (>99% CD4^+^ CD45RA^+^ CCR7^+^). γδ T-APCs were then irradiated at 12 Gy prior to coculture with allogeneic CD4+ T cells responder cells (1:10 ratio) for up to 9 d prior to analysis of surface phenotype and intracellular cytokine expression by flow cytometry.

### Cytokine staining

Cells were re-activated with PMA (10 ng/mL) and ionomycin (2 μM) in the presence of monensin (3 μM) for 4 h at 37°C, 5% CO_2_ prior to surface labelling. Re-activated cells were permeabilized using Leucoperm reagents (AbD Serotec, Oxford, UK), labelled with anti-cytokine mAb, and fixed in 4% paraformaldehyde for analysis by flow-cytometry.

### Flow-cytometry

Monoclonal antibodies were; PerCP-Cy5.5-conjugated CD3 (clone HIT3a), FITC-conjugated TCR Vδ2 (B6), Alexa Fluor®647-conjugated CD27 (O323), and CD103 (Ber-ACT8), PE/Cy7-conjugated CD45RA (HI100), and IFNγ (4S.B3), PE-conjugated integrin β7 (FIB504), CD69 (FN50), TNFα (MAb11), IL-17A (BL168), IL-10 (JES3-9D7) and BV421-CD49d (9F10), APC-conjugated CD86 (IT2.2), AlexaFluor®647-CD66b (G10F5), PECy7-CD69 (FN50), Pacific Blue-HLA-DR (L243), FITC-CD14 (HCD14) and PECy7-CD16 (B73.1) from BioLegend UK. PE/Cy7-conjugated anti-IL-22 (22URTI) was from eBioscience. APC-H7-conjugated anti-HLA-DR (L243) was from BD Biosciences. Labelled cells were acquired on a FACSCanto II flow-cytometer using FACSDiva 6.1.2 software (Becton Dickinson), and data were analyzed using WinList 9.0 (Verity Software House, Maine).

### Fungal species identification

Fungal species were isolated during *in vitro* culture of mucosal biopsy samples from HC donors and patients with CD. Initial fungal cultures were streaked onto BD ChromAgar *Candida* plates and identified using AuxaColor20 biochemical assays. Some strains underwent further confirmative identification by MALDI-TOF due to ambiguous results from standard profiling (performed by the Royal London Pathology Laboratory).

### Candidalysin

Candidalysin (CLYS) toxin (SIIGIIMGILGNIPQVIQIIMSIVKAFKGNK) was synthesised commercially (Peptide Synthetics, UK), solubilised in culture-grade water, and stored at 10 mg/mL at -20 °C.

### Candida filamentation assay

*Candida* isolates were cultured in YPD for 16-18 h at 30°C then washed 3 x with PBS, resuspended in complete medium, and transferred to a 37°C incubator. At the time points indicated, hyphal growth was stopped by fixation in 4% PFA for 15 min. Filament length was then determined using widefield microscopy (IN Cell Analyzer 2200, GE Healthcare) and analysis by ImageJ2 v2.3.0.

### Quantification of ECE1 gene expression

Epithelial cells were infected with *Candida* species/strains for 24 h (MOI 0.01). Exhausted culture medium was collected, and non-adherent fungi were pelleted by centrifugation. Adherent fungi were rinsed with ice-cold PBS, loosened with a cell scraper, and added to the pelleted sample. Fungi were washed with 1 mL ice-cold PBS, resuspended in a final volume of 500 µL, and added to a cryo-vial containing approximately 1/3 volume of acid-washed glass beads (Thistle Scientific, 0.5 mm) for 4 rounds of bead beating at 4.5 m/s^2^ for 30 s using a FastPrep-24 system (MP Bio). Lysates were removed and RNA was extracted using a MasterPure Yeast RNA Purification Kit (Lucigen) according to the manufacturer’s instructions. Removal of contaminating gDNA was performed according to kit instructions. For t = 0 h control samples, RNA was extracted from 500 µL of each YPD fungal culture in the absence of epithelial cells. cDNA was synthesized using a QuantiTect Reverse Transcription Kit (Qiagen) and 600 ng of RNA template. cDNA samples were then used for qPCR using HOT FIREPol EvaGreen qPCR Supermix (Solis BioDyne). Primers (universal forward and reverse for *ACT1*, species specific forward and reverse for *ECE1*) were used at a final concentration of 200 nM. qPCR amplifications were performed using a Rotor-gene system (Corbett). *ECE1* gene expression was calculated individually for each strain during culture on Caco-2 epithelial cells using the threshold cycle (ΔΔCT) method to calculate fold change (2^ ΔΔCT) against *ACT1* reference gene. Data shown represent fold change of expression relative to the SC5314 reference control (set as 1). The following oligonucleotide primers (5’-3’) were used to quantify *ECE1* gene expression. *C. albicans* SC5314 (Fw: ctttatcttctcaagctgc, Rev: caacaacagaatcaatatcttc), *C. dubliniensis* CD36 (Fw: gctgatcctgttgttgctgaacc, Rev: atggcatatcagcaatgacaccag), *C. tropicalis* MYA-3404 (Fw: gatgctgtcttagctggttctg, Rev: catctctcttaacaaggccagc), and universal ACT1 (Fw: ccaggtattgctgaacgtatgc, Rev: ggaccagattcgtcgtattcttg).

### Quantification of cellular damage

Caco-2 cells were challenged with fungi at MOI 0.01 for 24 h. A Cytox 96 non-radioactive cytotoxicity assay kit (Promega) was used according to the manufacturer’s instructions. Recombinant porcine lactate dehydrogenase (Sigma) was used to create a standard curve.

### Fungal sequencing

*C. albicans* genomic DNA was obtained using the spheroplasting guanidine extraction method to harvest high molecular weight DNA. *C. albicans* cells (10 mL) grown overnight on YPD (30°C, 200 rpm shaking) were harvested by centrifugation at 650 x g for 4 min and the supernatant was removed. The cell pellet was frozen at -20 °C for 30 min before resuspending in 0.5 mL spheroplasting buffer (36.43 g sorbitol plus 10 mL 1 M potassium phosphate buffer (pH 7.0), 4 mL 0.5 M EDTA (pH 7.5) and H_2_O to a volume of 200 mL (filter sterilized). Prior to use, 1% (v/v) 2-mercaptoethanol was added and incubated for 15 min at 37 °C. Spheroplasts were harvested by centrifugation at 750 x g for 4 min and the supernatant discarded. A 1.5mL volume of guanidine solution (21.5 g Guanidine-HCl plus 10 mL 0.5 M EDTA, 0.44 g NaCL, 0.25 mL of 10% sarkosyl and H_2_O to a volume of 50 mL, adjusted to pH 8.0) was added and the pellet gently resuspended. Pellets were incubated at 65 °C for 20 min before cooling on ice. Ethanol (1.5 mL of 100%) was added, samples inverted, and stored at -20 °C overnight. Pellets were treated with 0.5 mL RNase solution (10 x TE buffer [pH 8.0] plus 50 μg/mL RNase A [DNase free]) for 1 h at 37 °C. Proteinase-K solution (15 μL of 20 mg/mL in 20 mM CaCl2 10 mM Tris-HCl [pH 7.5]) was then added to tubes and incubated for a further 1 h at 65 °C. DNA was then extracted by adding 0.5 mL phenol:chloroform:isoamyl alcohol (Sigma-Aldrich, UK), inverting and pulsing in a microcentrifuge. The upper DNA-containing layer was transferred to a fresh tube and phenol:chloroform:isoamyl alcohol extraction was repeated. DNA was precipitated by adding 1/20 volume of 3M sodium acetate (pH 5.2) and 1/20 volume of 100% ethanol, microcentrifuging at full speed, then the ethanol was completely removed. DNA pellets were air-dried for 20 min before resuspending in 15 μL water. Extracted genomic DNA was sequenced using Illumina methodology in the Centre for Genome Enabled Medicine (University of Aberdeen).

### Bioinformatics

Read quality was first assessed using FastQC (v0.12.1) and low-quality bases and adapters were trimmed using TrimGalore (v0.6.5). Reads were then aligned to the reference genome using Bowtie2 (v2.4.5) and alignment metrics were gathered and analysed using Samtools (v1.10). Variants were called using the GATK HaplotypeCaller pipeline. In brief, aligned SAM files were sorted, converted to BAM files and indexed, then the HaplotypeCaller function was called on the processed BAM files. Variants called by the HaplotypeCaller were then hard filtered with the following recommended thresholds; “QD < 2.0”, “FS > 60.0”, “MQ < 40.0”, “SOR > 4.0”, “MQRankSum < -12.5”, “ReadPosRankSum < -8.0”. Base Quality Score

Recalibration was then performed (ApplyBQSR), using the filtered variant calls and the previously generated BAM file. Variants were then recalled using the HaplotypeCaller function. Finally, variants called using the recalibrated BAM file were filtered using the same thresholds as above to produce a final VCF file for each sample. VCF files for each sample were merged into a single cohort VCF file using bcftools (v1.19), then indexed and converted to a table using the GATK VariantsToTable function. To produce phylogeny trees, the cohort VCF file generated above was converted to a phylip file using the vcf2phylip package (v2.8). This phylip file was then processed using RAxML (v8.2.12) with GTR, optimization of substitution rates, and GAMMA model of rate heterogeneity (without bootstrapping). Finally, the GTRGAMMA model was used with 100 bootstraps to produce a “best tree”.

### Pulsed SILAC proteomics

The mass spectrometry proteomic data have been deposited to the ProteomeXchange Consortium via the PRIDE [1] partner repository with the dataset identifier PXD027433. Selected *C. albicans* strains were cultured overnight in YNB medium (supplemented with sucrose and ammonium sulphate) in either normal oxygen or hypoxic conditions (∼98% N_2_, ∼2% O_2_). The cultures were then switched to pSILAC medium (R and K-free RPMI medium supplemented with K8 and R6 amino acids). Cultures were maintained under these conditions for 3 h before the cells were collected for pSILAC proteomic analysis. Data were input into FungiDB (https://fungidb.org/fungidb/app/) and CGD Gene Ontology Term Finder (http://www.candidagenome.org/cgi-bin/GO/goTermFinder) to identify major biological pathways upregulated within the proteomic profiles of CA16 and CA47 under hypoxic or normal oxygen conditions. Enrichment analysis was performed using REVIGO^110^ web server (http://revigo.irb.hr/) to generate results for each strain/condition in two-level hierarchical format, from which the most significant gene ontology terms were rendered as pie charts using CirGO software^111^.

### *C. albicans* transformations

Transformations were performed as described in Gratacap *et al*.^76^. Briefly, strains were constructed by transforming CA16 and CA47 with the pENO1-iRFP-NATr plasmid which contains a codon-optimized version of the iRFP gene under the control of constitutive *ENO1* promoter, as well as a nourseothricin resistance (NATr) selection marker (pUC57 backbone; Genscript, Germany). The transformation was carried out using lithium acetate as previously published^112^, with nourseothricin resistance as an integration marker (100 μg/mL NAT, Werner Bioagents). Twenty colonies were selected and screened for fluorescence by flow cytometry (488/585 nm, FACScalibur, Becton Dickinson). A PCR check for integration was performed using the following primers to verify correct plasmid integration (1185bp): pENO1 FW: 5’tccttggctggcactgaactcg-3’ and iRFP REV: 5-5’atcacatgaagtcaaatcaacttttctagc-3’. Successful transformation was confirmed via fluorescence microscopy of fungal strains using a ZEISS Imager M2 upright microscope.

### Gut-on-a-chip infection assay

Intestine S1 chips were obtained from Emulate (Boston, MA). Chips were activated for ECM coating as per the manufacturer instructions. Chips were activated using ER-1 solution (Emulate) and exposed for 20 min under UV light (36 W, 365 nm), then rinsed with ER-2 solution followed by PBS (Gibco). Next, ECM coating solution (100 mg/mL matrigel [Corning] and 30 μg/mL rat tail collagen type 1 [Gibco] diluted in DMEM [Gibco]) was added and incubated for 4 h at 37°C 5% CO_2_. HUVEC cells were seeded at a concentration of 6 x 10^6^/mL in the bottom channel and the chips inverted for 2 h at 37°C 5% CO_2_. After cell attachment, chips were turned upright and washed with HUVEC medium supplemented with 1% Pen-Strep and 1% L-glutamine. Caco-2 cells were then seeded into the bottom channel at a concentration of 3 x 10^6^/mL and incubated for 2 h. Chips were gravity washed before remaining static overnight. Finally, chips were connected to the Emulate platform and maintained for the duration of culture using the following conditions: 37°C, 5% CO_2_, 30 μL/h flow, 10% lateral mechanical stretching at a frequency of 0.15 Hz. Cell culture medium was equilibrated and refreshed every 3 d.

At the start of the experiment timeline, the inlet reservoir for the top channel was filled with Caco-2 culture medium containing a known concentration of fungi (8 x 10^4^/mL). This was then flowed through the chip for 30 min at 400 µL/h to the top channel and 30 µL/h to the bottom channel with no stretch applied to synchronize the medium. This chip was imaged for the t = 0 time point to visualise fungal adhesion before incubating for a further 30 min at 30 µL/h with flow and stretch applied (10% and 0.15 Hz). The channels were then washed through with complete medium supplied via the inlet ports at 400 µL/h with no flow or stretch applied. Each channel was washed a further three times with PBS before the channels were fixed with 4% PFA for 20 min. After incubation, the channels were washed three times with PBS before ensuring that channels were filled with PBS and chips were submerged in PBS for storage at 4°C until subsequent imaging.

### Gut-on-a-chip immunofluorescence

Organ chip channels were washed through with PBS and then stained to visualise hyphal invasion and structural features. Each channel was filled with 100 µL blocking buffer and tips filled with buffer were left in the ports. Chips were then incubated either overnight at 4°C or for 2 h at room temperature in a humidified box. Tips were removed from ports and channels were washed three times with PBS. Primary antibodies were made up in dilution buffer and 100 µL of the mix was added to both channels and left overnight at 4°C in a humidified box. Primary antibodies used were 1:200 rabbit anti-ZO-1 (Abcam, EPR19945-296) and 1:200 mouse anti-E-cadherin (Abcam, HECD-1). After incubation, both channels were washed three times with PBS before 100 µL of secondary antibody mix was added and the chips left to incubate for 2 h in a humidified box at room temperature in the dark. Secondary antibodies used were 1:500 AF488 phalloidin, 1:500 AF568 goat anti-rabbit, 1:500 AF647 donkey anti-mouse (ThermoFisher) made up in dilution buffer. After incubation, both channels were washed through with PBS three times. Chips were submerged in PBS until ready for imaging, at which point they were mounted onto clear coverslips with ProlongFade and PBS. Imaging was performed using an inverted Zeiss 880 Laser Scanning Confocal Microscope with AiryScan and analysis was conducted on ImageJ (Version 2.3.0).

### Statistics

Statistical analyses were conducted using Prism 9 software (GraphPad, USA). Data normality was assessed using Anderson-Darling, D’Agostino & Pearson, Shapiro-Wilk, and Kolmogorov-Smirnov tests as indicated. For unpaired data, Mann Whitney test was used to compare two groups assuming unequal variance, while one-way ANOVA with a Kruskal-Wallis post-hoc test was employed for comparisons involving more than two groups. A paired t-test was used for two-group comparisons, and a repeated measures two-way ANOVA with Bonferroni’s multiple comparisons post-hoc test was applied for comparisons involving more than two groups. Statistical significance was defined as a p-value of <0.05.

## Acknowledgements

The authors wish to thank Dr Jan Soetaert and Dr Luke Gammon (Blizard Institute, QMUL) for technical support with confocal microscopy and high-content image analysis and processing, as well as Dr Mina Mincheva (Centre for Predictive *in vitro* Models, QMUL) for assistance with organ-on-a-chip sectioning and visualization. LM was supported by a Large Project Grant from Bart’s Charity (MGU0465). SC received an Academic Clinical Fellow support grant from Bart’s Charity (G-002461) and is supported by the Health Advances in Underrepresented Populations and Diseases (HARP) PhD programme (Wellcome Trust: 223500/Z/21/Z). JRN was supported by grants from the Wellcome Trust (214229/Z/18/Z) and Biotechnology and Biological Sciences Research Council (BBSRC: UKRI717). NEM was supported by a Career Development Award from the Medical Research Council (MRC: MR/R008302/1) and a Large Project Grant from Bart’s Charity (MGU0465).

## Author contributions

**Liya Mathew**: Conceptualization, Data curation, Formal analysis, Investigation, Methodology, Visualization, Writing – original draft, Writing – review & editing. **Sean Carlson**: Formal analysis, Investigation, Methodology, Visualization, Writing – original draft, Writing – review & editing. **Michael Savage**: Formal analysis, Investigation, Methodology, Visualization, Writing – original draft, Writing – review & editing. **Claire Pardieu**: Investigation, Methodology. **Megan O’Brien**: Investigation, Methodology. **Ellie Crawley**: Investigation, Methodology. **Paul Harrow**: Investigation, Methodology. **Ann-Kristen Kaune**: Data curation, Formal analysis, Investigation, Methodology, Writing – review & editing. **Edward Devlin**: Data curation, Formal analysis, Investigation, Methodology, Writing – review & editing. **Emily L. Priest**: Data curation, Formal analysis, Investigation, Methodology, Visualization, Writing – review & editing. **Marilena Crescente**: Investigation, Methodology, Writing – review & editing. **Kimberley Martinod**: Investigation, Methodology, Writing – review & editing. **James Boot**: Data curation, Formal analysis, Investigation, Methodology. **Marco Gasparetto**: Investigation, Methodology, Resources, Writing – review & editing. **Klaartje Kok**: Investigation, Methodology, Resources, Writing – review & editing. **James O. Lindsay**: Funding acquisition, Investigation, Methodology, Resources, Writing – review & editing. **Jonathan P. Richardson**: Formal analysis, Investigation, Methodology, Resources, Writing – review & editing. **Julian R. Naglik**: Formal analysis, Investigation, Methodology, Resources, Supervision, Writing – review & editing. **Andrew J. Stagg**: Funding acquisition, Investigation, Methodology, Writing – review & editing. **Matthias Eberl**: Conceptualization, Formal analysis, Funding acquisition, Investigation, Methodology, Resources, Writing – review & editing. **Siu Kwan Sze**: Data curation, Formal analysis, Investigation, Methodology, Resources, Validation, Visualization, Writing – review & editing. **Carol A. Munro**: Conceptualization, Formal analysis, Funding acquisition, Investigation, Methodology, Resources, Writing – review & editing. **Neil E. McCarthy**: Conceptualization, Data curation, Formal analysis, Funding acquisition, Investigation, Methodology, Project administration, Resources, Supervision, Visualization, Writing – original draft, Writing – review & editing.

