## Supplementary figures and images for "Bacterial suppression of intestinal fungi via activation of human gut γδ T-cells"

### Supplementary Figure 1

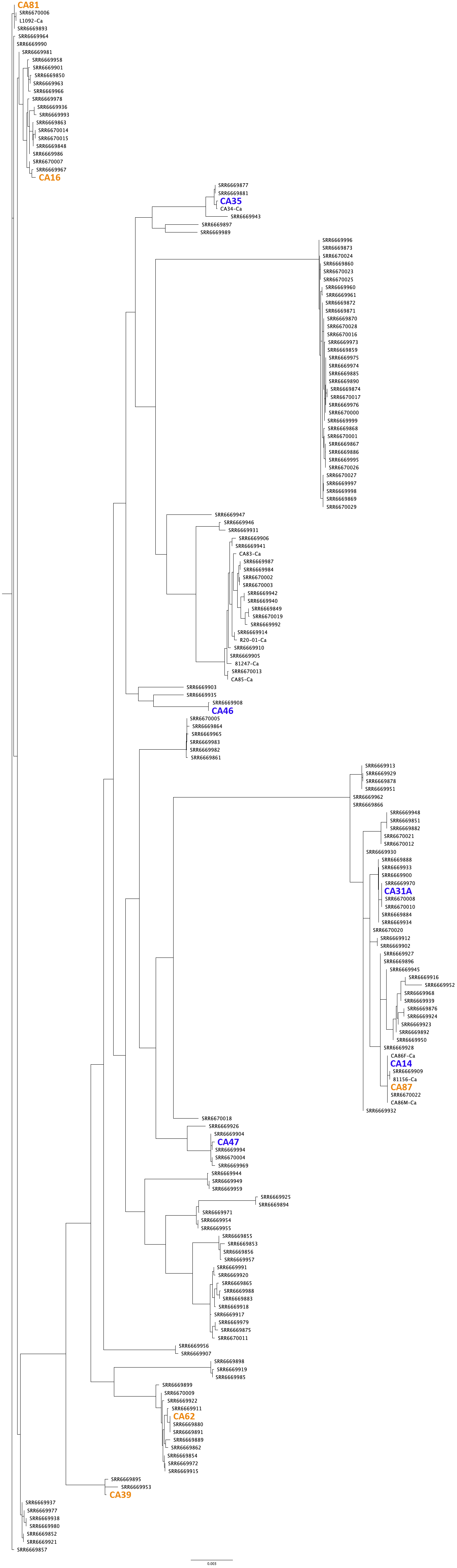

### Supplementary Figure 2

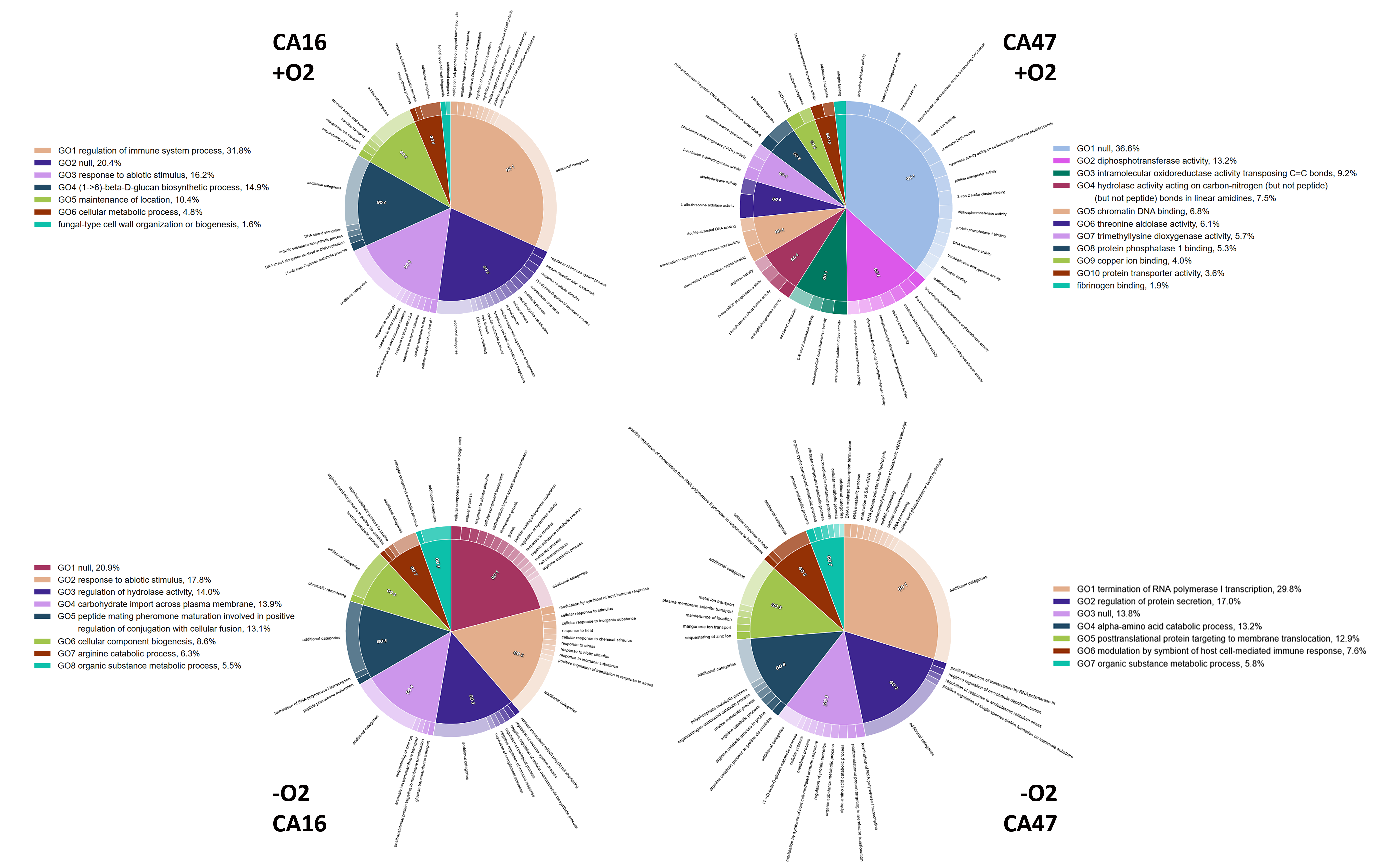
